# Blockiness as Determinant of Liquid Phase Formation by Polypeptide-Like Polymers in Stickers-Spacers Paradigm

**DOI:** 10.64898/2026.10.05.756799

**Authors:** Egor O. Vasilenko, Dmitry S. Belous, Dmitry N. Ivankov

## Abstract

Liquid–liquid phase separation (LLPS) driven by intrinsically disordered protein regions underlies the formation of membraneless organelles. The role of these structures in cell biology is an active area of research, yet their physical understanding remains less developed than that of protein assemblies with well-defined, rigid tertiary structures. The amino acid composition of disordered regions, particularly the abundance of charged residues, is known to modulate phase behavior. These factors govern accessible conformations, the assembly of compact sticker clusters, and the porosity of the resulting polymer networks. In contrast, the spatial distribution of side-chain volumes remains a comparatively understudied sequence determinant. Here, we employ a model that captures the patterning of side-chain volumes and interaction strengths without specifying chemical identity to show how sequence blockiness modulates the phase-nucleation propensity, connectivity, and compactness of polypeptide-like polymer assemblies. Using a library of polymers with identical composition but permuted residue arrangements, we show that clustering of bulky residues generates cooperative sticker domains and identify the threshold blockiness required to stabilize polymer networks. Coarse-grained simulations and mean-field theory reveal that this effect arises from an interplay among steric repulsion, solvent entropy, and higher-valency interactions, whereby blocky patterns promote the formation of multi-node percolated networks. Segregation of stickers and spacers is therefore one of the sequence parameters governing LLPS which may be useful for interpreting the evolution of disordered protein sequences, high-throughput proteome screening, and engineering of stimuli-responsive biomaterials. These rules may be exploited to predetermine condensate stability, viscoelasticity, and molecular exchange rates without altering chemical functionality.

## Introduction

Liquid–liquid phase separation (LLPS) underlies the formation of membraneless organelles (MLOs) [Hirose 2022, Posey 2018, Thakur 2025]. Such phase transitions occur when proteins reach a saturation concentration, at which the solution demixes into dense and dilute phases without a surrounding lipid membrane [Posey 2018]. Consequently, proteins and nucleic acids continuously exchange between the condensate and the surrounding medium. MLOs are dynamic compartments that assemble rapidly on demand, enabling cells to adapt fast to changes in their local environment. For example, the nucleolus concentrates ribosomal proteins and RNA to facilitate ribosome biogenesis [Chen 2001]; stress granules transiently accumulate untranslated mRNAs and translation factors during stress [Franzmann 2019]; and P-bodies serve as sites for RNA storage and degradation [Jain 2013]. By locally concentrating specific proteins, particularly transcription factors, together with nucleic acids, MLOs facilitate specific biochemical reactions, thereby providing spatial and temporal control over gene expression and other cellular processes [Hirose 2022, Driver 2025]. Elucidating the molecular principles that govern LLPS is therefore essential for understanding how eukaryotic cells achieve such precise regulation of fundamental biological functions [Hirose 2022, Wang 2021]. The proteins that nucleate and sustain these condensates predominantly contain intrinsically disordered regions (IDRs), which lack stable secondary and tertiary structures [Wright 1999, Trivedi 2022]. Consequently, IDRs engage in numerous weak, transient contacts that collectively give rise to multivalent interactions, the primary driving force for LLPS [Franzmann 2019, Martin 2020]. Because IDRs do not conform to the conventional structure-function paradigm [Wright 1999], identifying the sequence features that determine their phase-separation behavior remains a central challenge [Ruff 2022].

A common approach is to describe IDR patterns and functions in terms of molecular grammars [Ruff 2025, Ruff 2022], which aim to identify the physicochemical properties governing phase-separation behavior directly from the amino acid sequence rather than from stable secondary or tertiary structure [Ruff 2022]. Proteins capable of independently driving LLPS while simultaneously serving as a matrix for other proteins or nucleic acids during condensate assembly have been designated by several terms, including scaffold, driver, and PS-self [You 2026]. Their interaction partners, termed clients or PS-other [You 2026], partition into this network but cannot independently nucleate a protein-rich phase. Throughout this work, we adopt the driver-client terminology to distinguish proteins explicitly by their functional role in condensate formation. Clients can shift the saturation concentration and alter the material properties of the condensate. The resulting condensates can adopt a range of material states, from liquids to gels and fibers [Franzmann 2019]. The physical basis of LLPS is commonly described by the stickers-and-spacers model [Mittag 2022, Martin 2020]. In this representation, driver proteins behave as self-assembling polymers composed of interacting sticker regions separated by flexible spacers. Stickers mediate noncovalent intermolecular interactions, including aromatic, electrostatic, hydrophobic, and cation-π contacts, whereas spacers primarily maintain chain flexibility and solubility [Franzmann 2019, Martin 2020]. Above a critical concentration, these crosslinks form a system-spanning network that undergoes phase separation [Posey 2018]. The stickers-and-spacers paradigm implies that both amino acid composition and the spatial arrangement of residues along the sequence influence phase behavior [Cohan 2022, Ji 2026]. Indeed, previous studies have demonstrated that charge segregation, or charge blockiness, strongly modulates LLPS propensity [Yamazaki 2022, Bianchi 2024, Sundaravadivelu Devarajan 2024]. Proteins with more pronounced blockiness in their charge and aromatic residue distributions phase-separate at lower concentrations and form denser condensates than those with uniformly distributed charges [Martin 2020, Sundaravadivelu Devarajan 2024]. Unambiguous roles of protein sequence regions are also difficult to establish: molecular grammars can be computed using multiple definitions, and phase behavior is often assessed qualitatively without molecular-level insight. To obtain a general understanding, one can combine grammar prediction methods with molecular dynamics simulations to evaluate the resulting predictions in a physical model. The physical consequences of residue composition and distribution can be captured using coarse-grained simulations [Zhang 2022, Rizuan 2026]. For example, the recently published CASPULE tool [Chattaraj 2026] employs a one-bead-per-residue polypeptide model. The growing number of such tools reflects the demand for physics-based, high-throughput sequence screening and the evaluation of patterning rules. More detailed models [Zhang 2022] employing multiple beads per residue also account for hydrogen bonds between backbone atoms, which are important for amyloid aggregation, but can be omitted for LLPS research. This omission is not a critical limitation for LLPS simulations, which are rather governed with non-specific transient interaction between side chains, and topological restrictions like entanglement of loops.

To focus on simulations of polypeptide condensates, here we developed and employed the LazyPhase tool (Fig. 1), which relies on a two-bead-per-residue representation [Szała-Mendyk 2022] that captures sequence features dependent on Ramachandran angles. To incorporate flexibility, volume, and interaction strength, we adopted a generalized sticker side-chain model in which sticker residues exhibit extended interaction ranges while occupying larger excluded volumes. To justify this framework, we addressed the question of whether block-like organization of different physicochemical properties commits to LLPS. Using sliding-window analysis of known LLPS-forming sequences, we found that residues with similar solvent-accessible surface area (SASA) values are likewise organized into contiguous blocks in driver proteins, whereas client proteins exhibit a more uniform distribution. To examine the consequences of this organization, we performed coarse-grained molecular dynamics simulations of polymers with varying degrees of blockiness [Zhang 2022, Chattaraj 2026, Szała-Mendyk 2022, Rizuan 2026]. Our results demonstrate that increasing the blockiness of generalized stickers is sufficient to promote aggregation while preserving the chemical composition of the polymer, leading to the formation of more compact clusters.

**Fig. 1.**
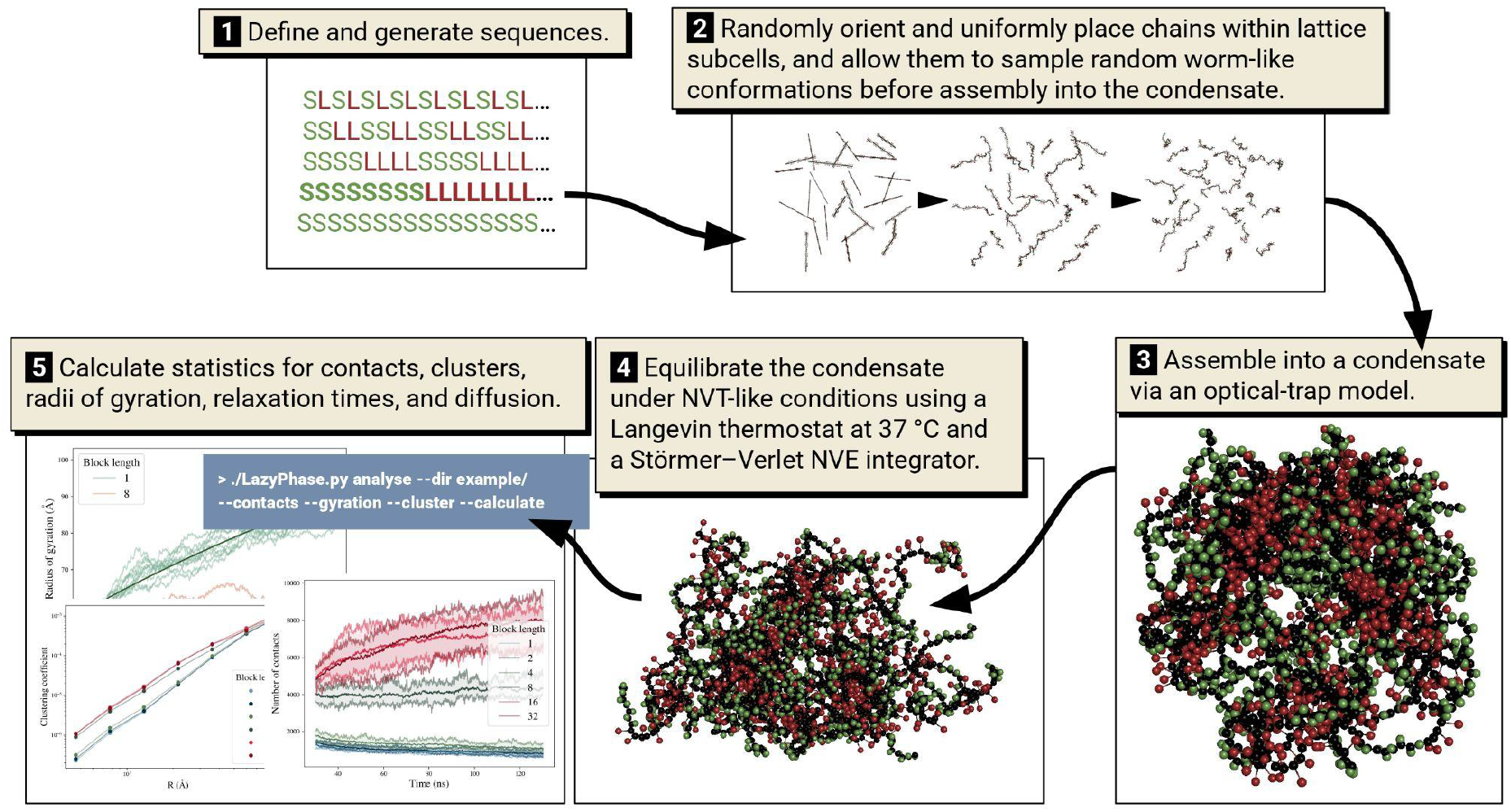
High-throughput computational workflow of LazyPhase, from blockwise sequence generation through trapped condensate fixation to analysis of polymer network properties.

## Methods

### Extraction and filtering of LLPS data

To analyze residue distributions in LLPS-forming sequences, we used the DrLLPS [Ning 2020] and PhaSepDB 3.0 [You 2026] databases, which contain annotated driver and client sequences (Supplementary Data), and screened 92,709 client and 2,808 driver sequences. To assess the statistical significance of differences in heterogeneity between these two sequence classes, we applied the Mann–Whitney U test to heterogeneity distributions. We used sliding-window scoring analogous to the NARDINI approach [Cohan 2023] (Fig. 2).

**Fig. 2.**
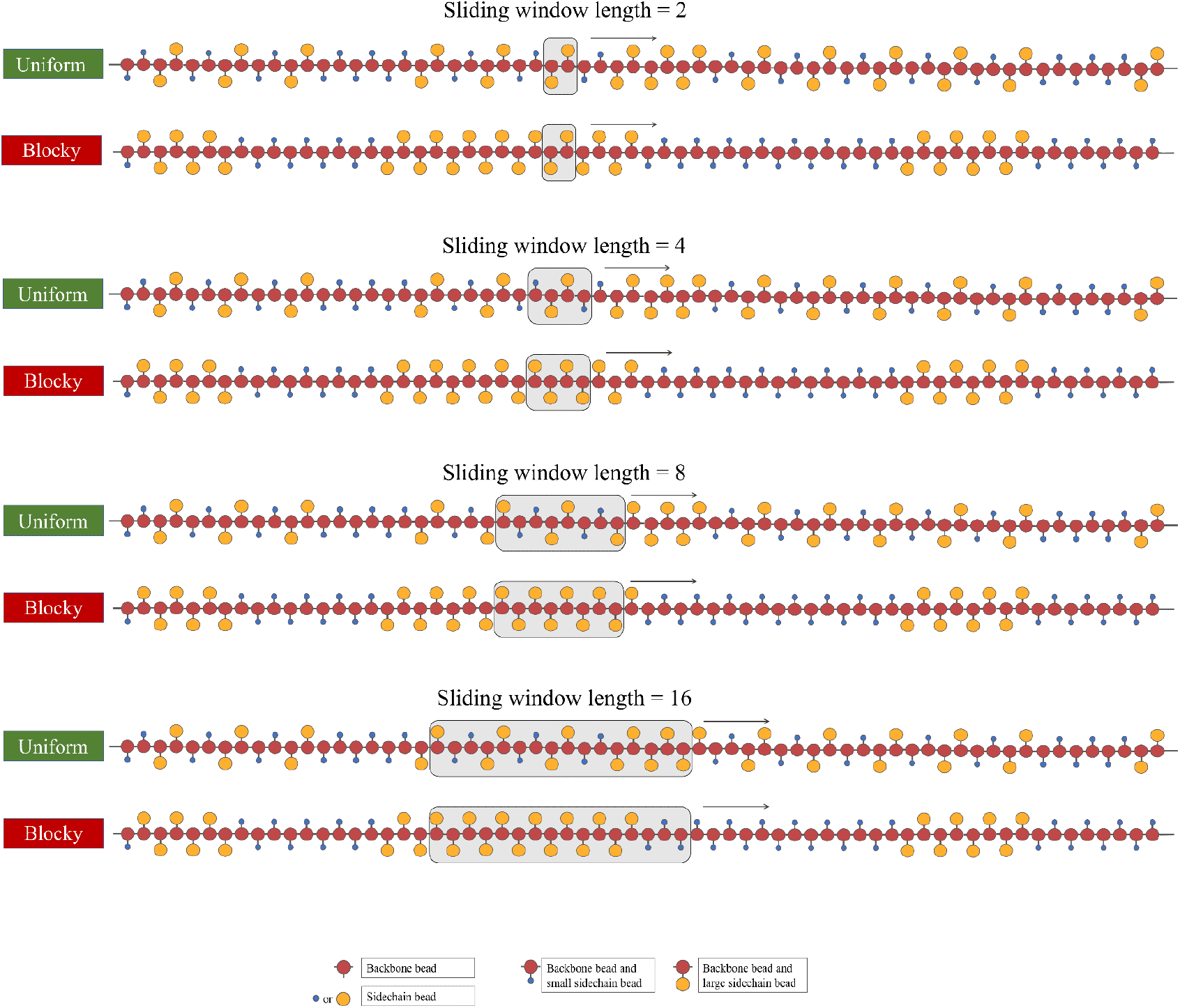
Schematic depiction of the sliding-window algorithm applied to more uniform and more blocky sequences, using window sizes of 2, 4, 8 and 16 residues.

### Calculation of heterogeneity scores

For a window of length *w* starting at position *i*, the score for property *p* is defined as:

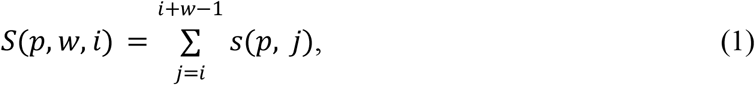

where *s(p, j)* is the value of property *p* at position *j*. The normalized heterogeneity *a(p, w)* for property *p* and window length *w* was calculated as:

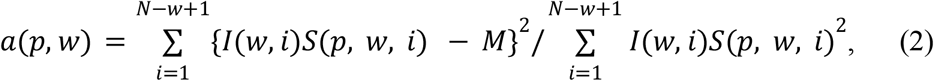

where *I(w, i)* is an indicator function equal to one if the mean IDR propensity in this region predicted by AIUPred [Erdős 20024] exceeds 0.5, and zero otherwise, and 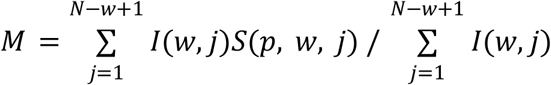 is the mean score over all windows in the sequence. The heterogeneity defined in Eq. (2) is equivalent to the variance of the window scores normalized by the sum of squares:

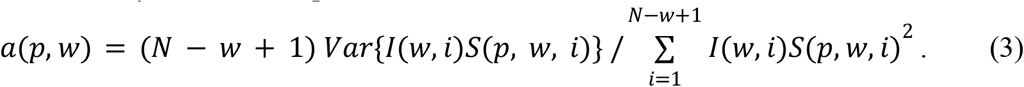

A sliding window of length *w* yields a set of scores along the sequence, {S_1_, S_2_, S_3_, …, S_N-w+1_}. For every client and driver sequence, we calculated the squared deviations for each such set and normalized them to 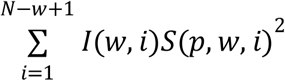. This yielded two sets of normalized deviations, one for clients and one for drivers. We then performed a Mann–Whitney U test to determine whether there was a statistically significant difference between client and driver heterogeneity. The block lengths were chosen to be at least an order of magnitude shorter than the average sequence lengths of both drivers and clients (Fig. S1).

### Mean-field theory

It is well established that heterogeneity facilitates the nucleation of new phases, particularly protein folding [Lequerica-Mateos 2024]. In addition to the likely energetic contribution of higher-valence pairings, a concomitant reduction in entropic penalty through clustering of interacting monomers can be expected. Here, we generalize the reduced-space effect [Choi 2020] to bound stickers of blockiness *b*, assuming that the contacts between residues within a cluster can permute:

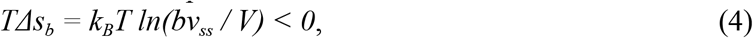

yielding a rough estimate of the entropy gain arising from segregation of sticky residues along the polymer relative to a uniform distribution:

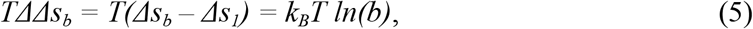

corresponding to an entropic advantage of approximately 2 *k*_*B*_*T*.

The dynamics of the radius of gyration were approximated using the three-dimensional diffusion equation [Wedemeier 2009]:

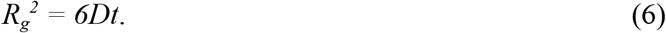

### Molecular dynamics (MD)

We developed a new tool, LazyPhase, designed to explore sequence effects on polypeptide self-assembly. In each simulation, 27 polymers were placed uniformly on a lattice and rotated randomly around their geometric centers. The polymers were allowed to sample random conformations in implicit solvent for 10 ns, then to assemble into and maintain a trapped condensate for 10 ns, and finally to equilibrate for 110 ns under NVT production conditions. All simulations were performed with LAMMPS 2025 [Thompson 2022]. Systems were prepared using Moltemplate [Jewett 2021]. NVT-like conditions were approximated using a Langevin thermostat at 37 °C (310.15 K) combined with velocity-Verlet integration. Polypeptides were represented with backbone and side-chain beads, following the model of Szała-Mendyk et al. [Szała-Mendyk 2022] (Fig. 3).

**Fig. 3.**
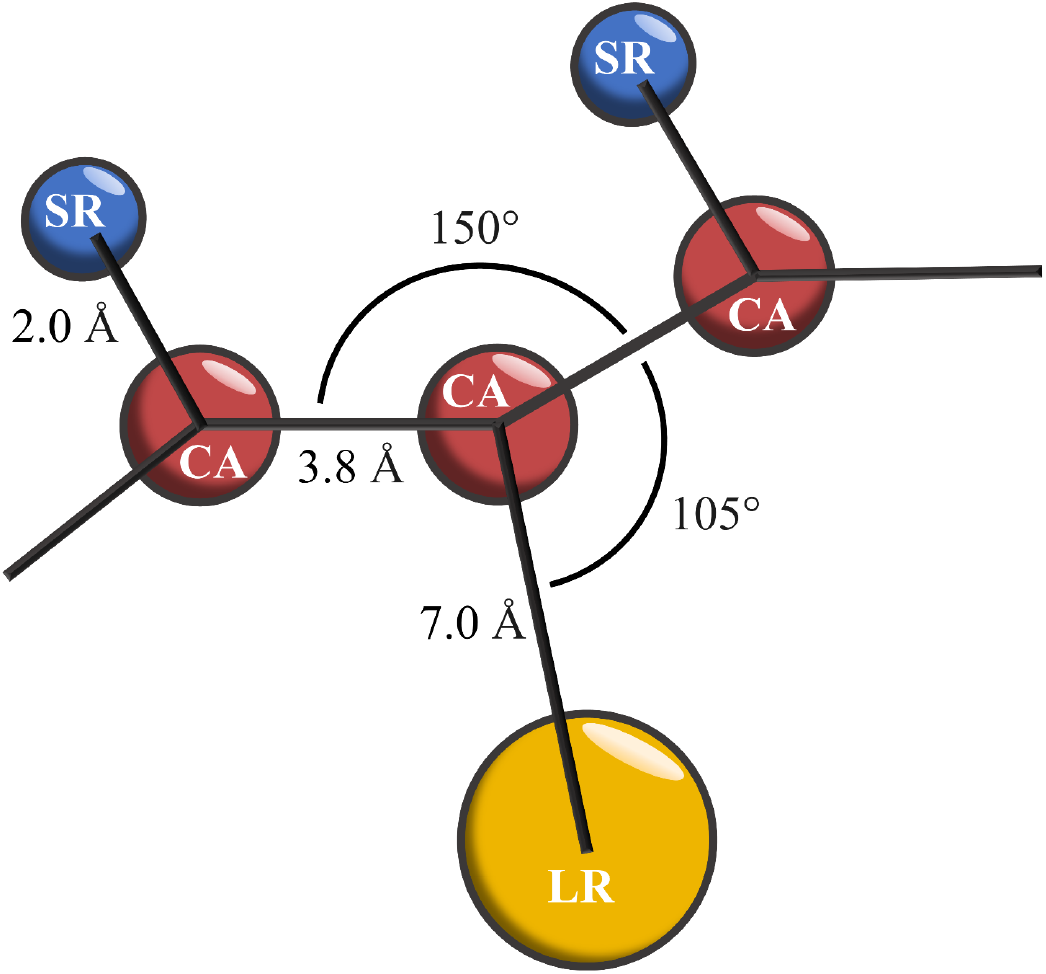
Schematic depiction of a three-residue segment in the coarse-grained two-bead-per-residue model used in this study.

Each polymer with the desired blockiness was generated using a Python script from the LazyPhase package (the link is provided at the end of the manuscript), which allows the user to choose a power of two that defines the block length, the polymer length, the number of polymers, and the approximate initial distance between polymers.

After placement and random rotation about their geometric centers and subsequent conformation sampling, the polymers were dragged to the center of the box. We employed an optical-trap model at the center of the simulation cell to eliminate the nucleation lag time and focus on prenucleus stability and transitions to nuclei; methods such as umbrella sampling have previously been used for polypeptide oligomerization [Davis 2010]. A similar approach was used to introduce a condensate into the simulation [Hunt 2016]. The purpose is to assemble the polymers into a compact state and observe the structures into which they reorganize over time.

All covalent bonds were modeled using a harmonic potential with stretching stiffness 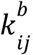, equilibrium bead-center distance 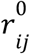, and bond energy 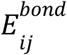 as a function of distance *r* _*ij*_ :

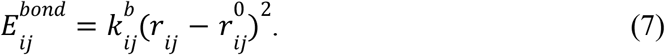

Angle bending between three consecutive covalently bonded beads contributes to the energy according to a cosine-squared potential, with 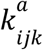 being the bending stiffness and θ_*ijk*_ and 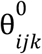 being the instantaneous and equilibrium angles, respectively, for beads with indices *i, j*, and *k*:

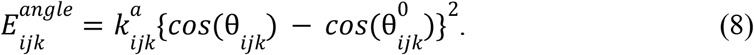

Nonbonded interactions between beads were described by the standard 12–6 Lennard-Jones potential, with σ_*ij*_ being the zero-crossing distance for beads *i* and *j*, and 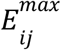 being the maximum absolute value of the interaction potential; this potential was not applied to covalently bonded beads:

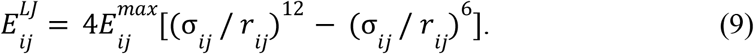

All parameter values are listed in Table I. The calculations of trajectories were repeated ten times for each sequence using different random seeds.

**Table I.** Model parameters used in all simulations.

| Bead type | Mass (g / mol) | Role |
| --- | --- | --- |
| CA (Cα) | 13 | Backbone α-carbon |
| SR (Small Residue) | 10 | Small sidechain |
| LR (Large Residue) | 50 | Large sidechain |

Nonbonded 12–6 Lennard-Jones interactions: interaction energies, zero-crossing distances, and cutoff distances.
| Pair | | $\epsilon$ (kcal / mol) | $\sigma$ (Å) | $r_{cut}$ (Å) |
| --- | --- | --- | --- | --- |
| CA | CA | 1.0 | 4 | 4 |
| SR | SR | 0.5 | 2 | 3 |
| LR | LR | 1.0 | 4 | 12 |
| CA | SR | 1.0 | 3 | 3 |
| CA | LR | 1.0 | 4 | 5 |
| SR | LR | 0.5 | 3 | 4 |

Harmonic bond potential; $k = 100 \text{ kcal} \cdot \text{mol}^{-1} \cdot \text{\AA}^{-2}$ .
| Bond type | Atoms | | $r_0$ (Å) |
| --- | --- | --- | --- |
| Backbone | CA | CA | 3.8 |
| Sidechain S | CA | SR | 2.0 |
| Sidechain L | CA | LR | 7.0 |

Equilibrium angles for the cosine-squared potential; $k = 20 \text{ kcal} \cdot \text{mol}^{-1} \cdot \text{rad}^{-2}$ .
| Angle type | Atoms | $\theta_0$ (°) |
| --- | --- | --- |
| Backbone | CA–CA–CA | 150 |
| Sidechain | CA–CA–*R | 105 |

| Free solution | Trapped | Relaxation | Main simulation |
| --- | --- | --- | --- |
| 10 | 10 | 10 | 100 |

### Analysis of MD trajectories

Radii of gyration were calculated using the canonical definition, where *R*_*COM*_ is the center of mass, with equal masses assigned to all beads, and *R*_*i*_ is the position vector of the *i*-th bead:

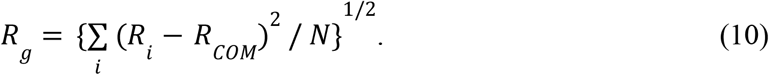

Two beads were considered to be in contact when the center-to-center distance fell within the range where the pairwise Lennard-Jones interaction energy is negative, excluding noninteracting pairs separated beyond the interaction cutoff or pairs in the overlap regions of steeply rising energy penalty:

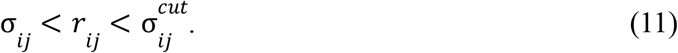

Clustering coefficients were calculated using the standard definition employed in hierarchical cluster analysis, fractal-dimension estimation, and investigations of system scaling. Two particles were assigned to the same cluster whenever their separation fell below the interaction cutoff, thereby defining a connected component in the same way as a contact in the contact statistics:

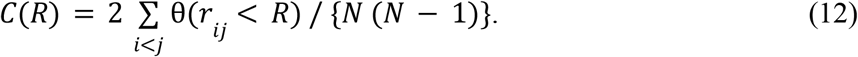

We modified this definition to represent intermolecular interactions exclusively. This correction was introduced to eliminate the contribution of intramolecular clustering in sequences with higher blockiness:

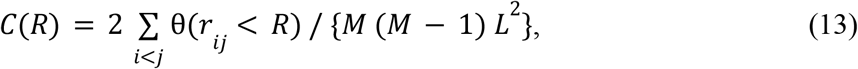

where θ(*x*) = 1 if condition *x* is satisfied, *M* denotes the number of molecules and *L* is the number of beads of the types of interest in a single molecule.

To accelerate the search for neighboring bead pairs, we used the k-d tree algorithm. To calculate the clustering coefficient at distance *R*, we constructed k-d trees and searched them with threshold *R*; the same acceleration technique was employed to identify contacts within the cutoff distance 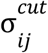.

## Results and Discussion

### Scale dependence of blockiness

Firstly, we compared the blockiness of driver and client sequences from DrLLPS [Ning 2020] and PhaSepDB 3.0 [You 2026] (Methods). We found that the driver sequences are generally more blocky than client sequences, and this is especially observed at larger scales. That means, the heterogeneity advantage of drivers over clients becomes more pronounced at longer window sizes. This observation motivates modeling driver sequences as blocky copolymers. Takakuwa et al. [Takakuwa 2026] reported that glutamine-rich regions facilitate MLO formation. In our calculations, drivers exhibit heterogeneity in glutamine comparable to or less than that of clients at the 2–8-residue scale but, in contrast, exceed at the 16-residue scale (Fig. 4). The distribution of general physicochemical properties, volume and flexibility, along the polypeptide chain, exhibits statistically significant differences between more segregated phase-nucleating driver sequences and phase-joining client sequences, motivating to introduce coarse molecule models incorporating side chain sizes. As shown in Fig. 4, drivers have higher heterogeneity scores of residue’s volume at every window length. In addition, distributions of histidine and leucine also demonstrate higher segregation in the drivers for lengths 8 and 16, which agrees with overall drivers prevalence in hydrophobic blockiness. Moreover, at a window length of 16 residues, drivers exhibited greater segregation across all properties, with the highest significance for the four charged residues: glutamate, aspartate, lysine, arginine, and for polarity.

**Fig. 4.**
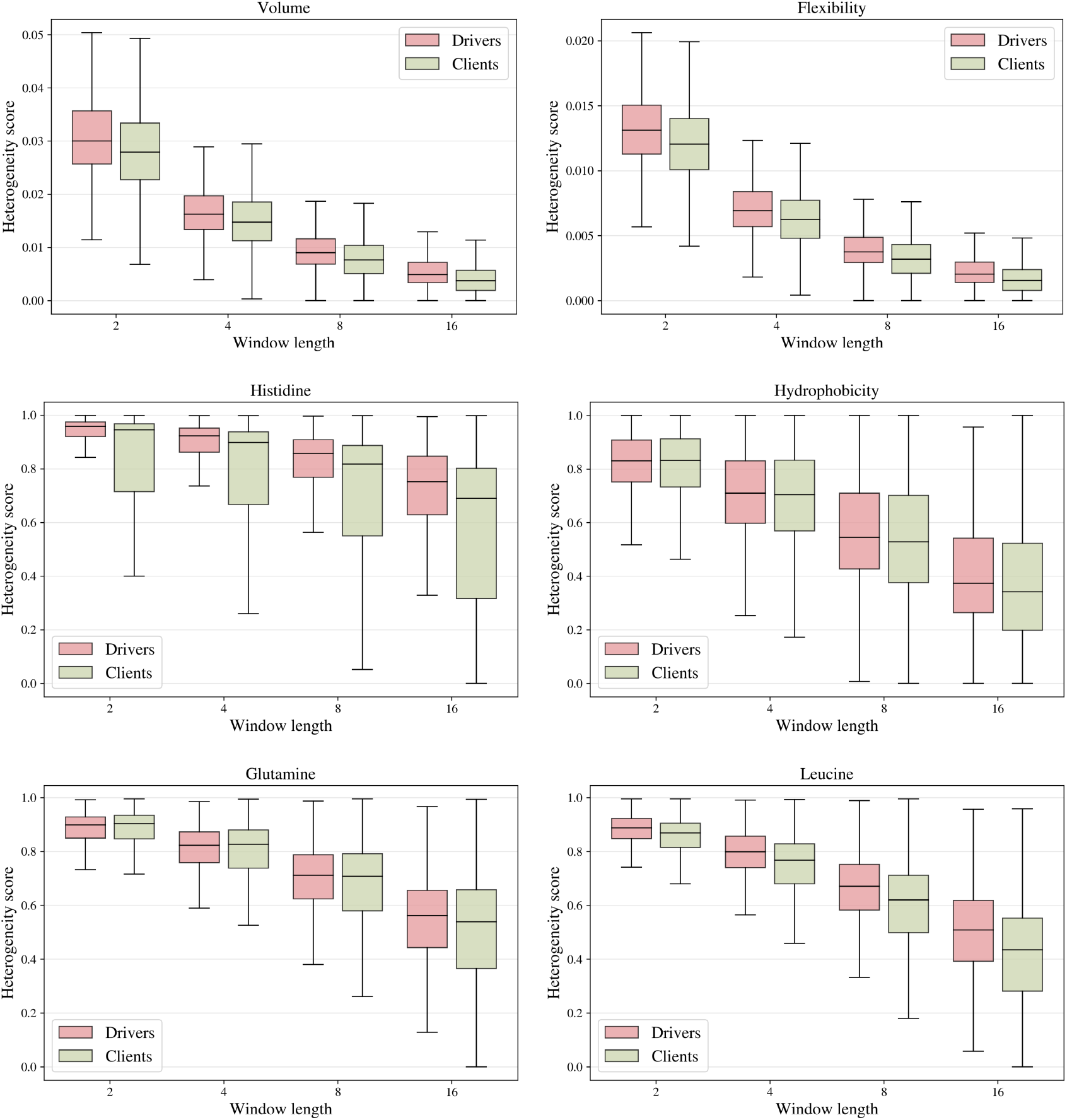
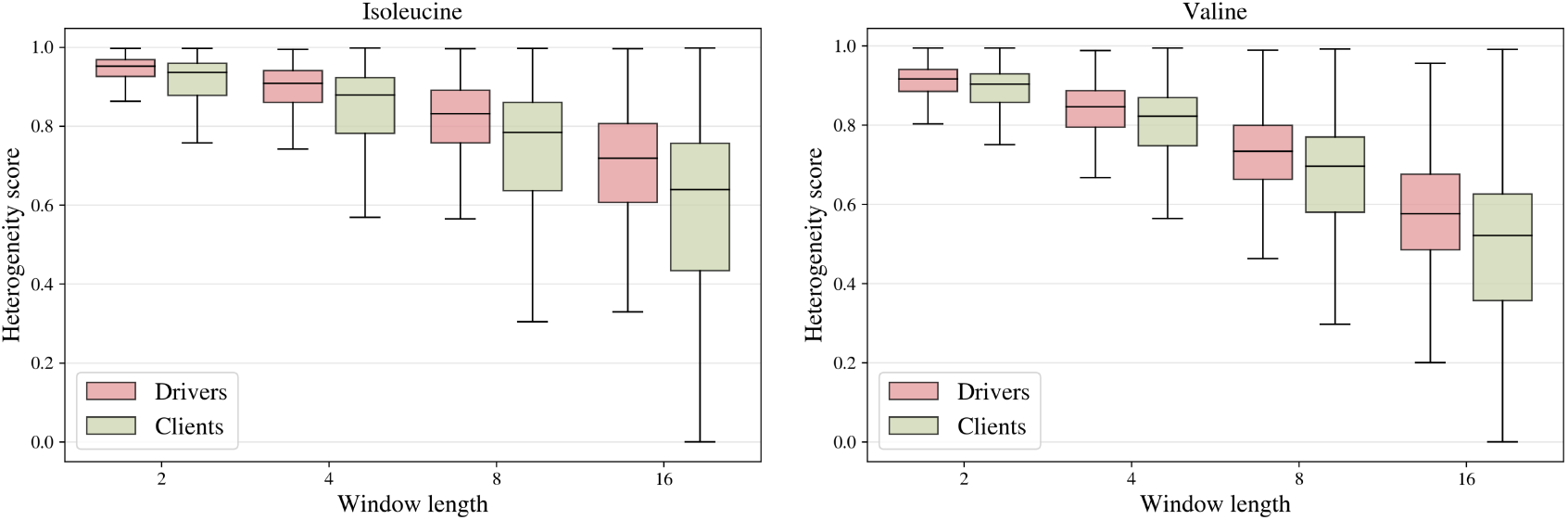
Normalized heterogeneities of several sequence properties computed using sliding windows of varying lengths. For a window length of 16 residues, all p-values are below 10^−4^.

### Contact statistics

The spatial arrangement of stickers along the polymer sequence and their degree of clustering are critical determinants of network formation. In Fig. 5, we show the final states of MD trajectories, which have different topologies depending on blockiness. Based on Fig. 6, we suggest that a threshold level of blockiness for 64-residue long polymers in our model exists between 4 and 8 residues per block, above which sufficient sticker length enables the formation of multiple nucleation cores and interconnected networks. There is a noticeable tendency: more blocky polymers tend to increase the number of contacts. Fig. 6 shows the evolution of the number of contacts in each type of polymer. Polymers with block lengths of 4–8 residues reach a plateau, whereas polymers with uniform distribution show a gradual decrease in the number of contacts, so the percolated network disintegrates. Clustered sticker blocks act as multivalent interaction sites, allowing multiple chains to simultaneously engage with the same region and promoting the formation of numerous interchain contacts, as is shown in Fig. 6. This reduces the entropic cost of crosslink formation and facilitates the emergence of percolated networks. The resulting networks remained highly dynamic, with transient crosslinks continuously forming and breaking, thereby generating an open porous architecture capable of accommodating client molecules of appropriate size and affinity. In contrast, uniformly distributed stickers form smaller and more isolated interaction regions, resulting in fewer simultaneous contacts between chains and requiring higher polymer concentrations to achieve phase separation.

**Fig. 5.**
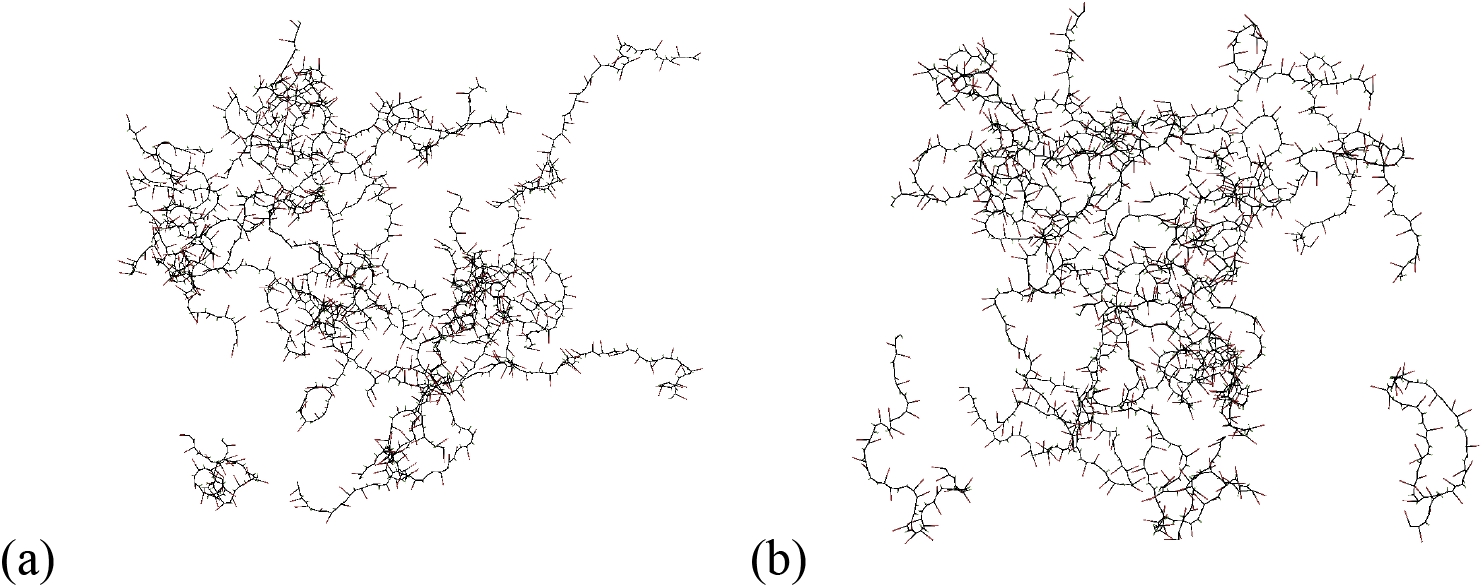
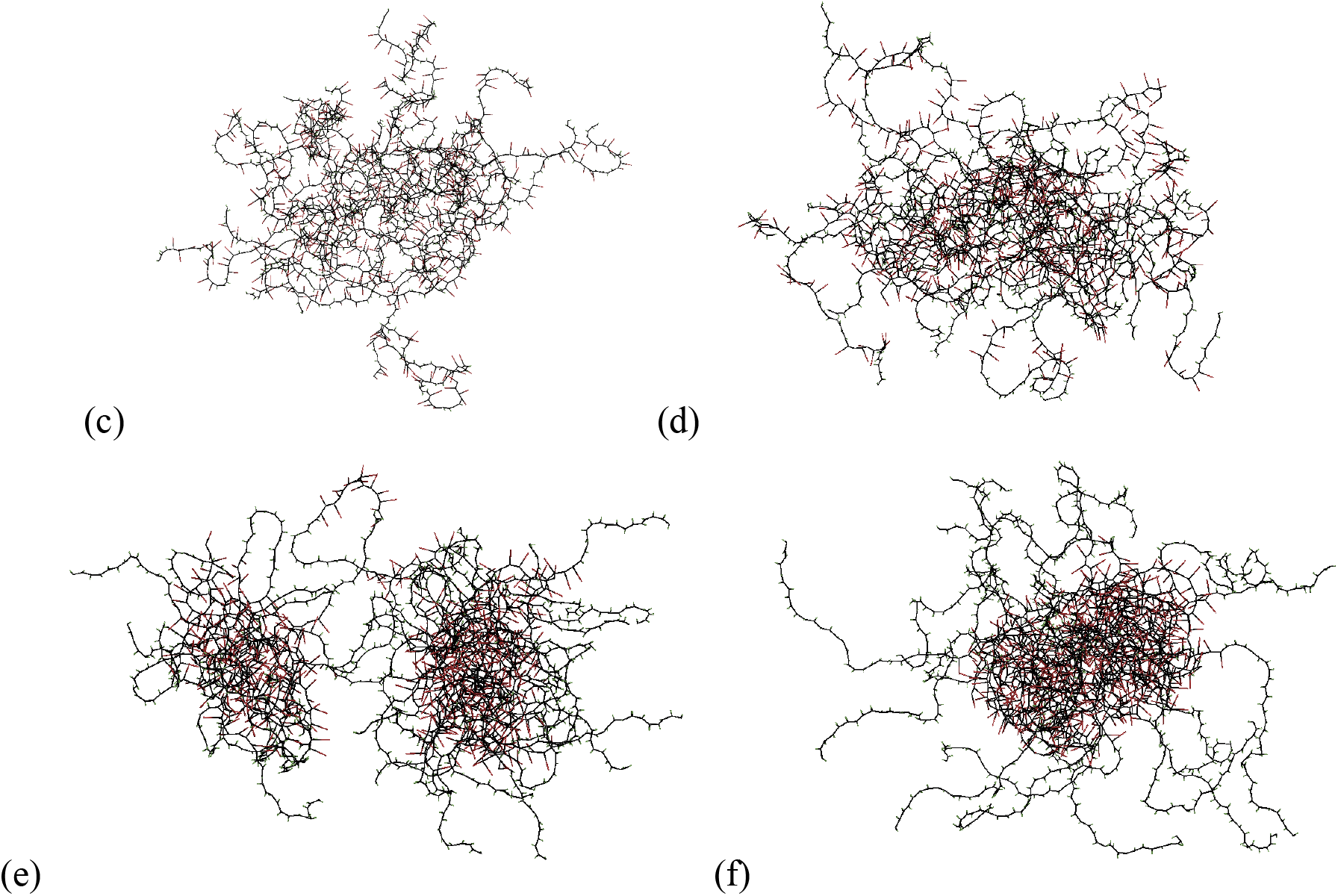
Example final states obtained for polymer networks of varying block lengths (a) 1; (b) 2; (c) 4; (d) 8; (e) 16; (f) 32.

**Fig. 6.**
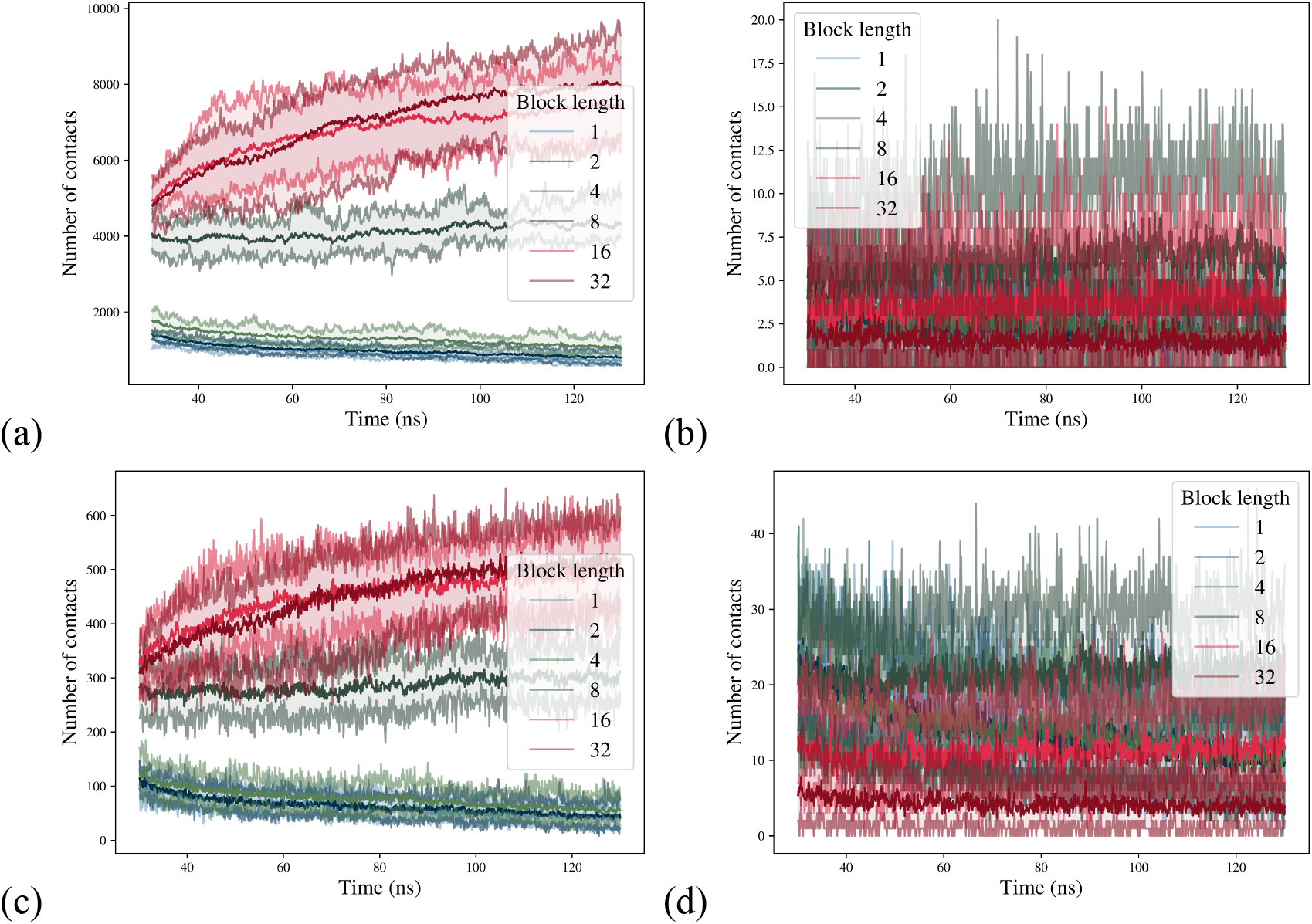
Time evolution of contact statistics, starting from trap force removal at 30 ns of simulation time. Pairs of bead types for which the contacts were calculated: (a) large-large; small-small; (c) backbone-large; (d) small-large.

### Radii of gyration

Client-like sequences assemble into less compact clusters than driver-like sequences do. As with other compactness measures, the radius of gyration of proteins correlates significantly with both folding rate and sequence length [Lobanov 2008, Ivankov 2009]; accordingly, we examined its behavior across different degrees of blockiness. This metric itself is not enough to judge, because increasing radius of gyration can indicate both diffusional sparsening of an unstable cluster and formation of stable brush-like rods. Coarse-grained simulations revealed a gradual transition in aggregate morphology with increasing residue blockiness (**Fig. 7**). Polymers with uniformly distributed large residues assembled into flexible percolated networks. The large radius of gyration shows the spread of the network relative to the common center of mass, which may indicate not only loss of condensate stability, but also formation of a micelle. Increasing the block length to 8–16 residues produced networks containing multiple compact nucleation cores connected by flexible strands. The lower radius of gyration indicates a more compact organization of the aggregate. Finally, polymers with the largest blocks, approximately half of the whole polymer, collapsed into a single dense aggregate composed of one large core surrounded by protruding spacer-rich tails, indicating a finite-size effect which limits the functional size of a block. In some simulations, the radius of gyration increased due to these stretching polymer fragments protruding from the main aggregate. We also note that the *R*_*g*_ dynamics of separate trajectories are highly stepwise, and we applied the averaged approximations by diffusion equation. Starting from the length of block 8, in some simulations we observed that artificially assembled nucleus relaxes and then rearranges into a more stable structure, as initially increasing radius of gyration eventually starts to decrease.

**Fig. 7.**
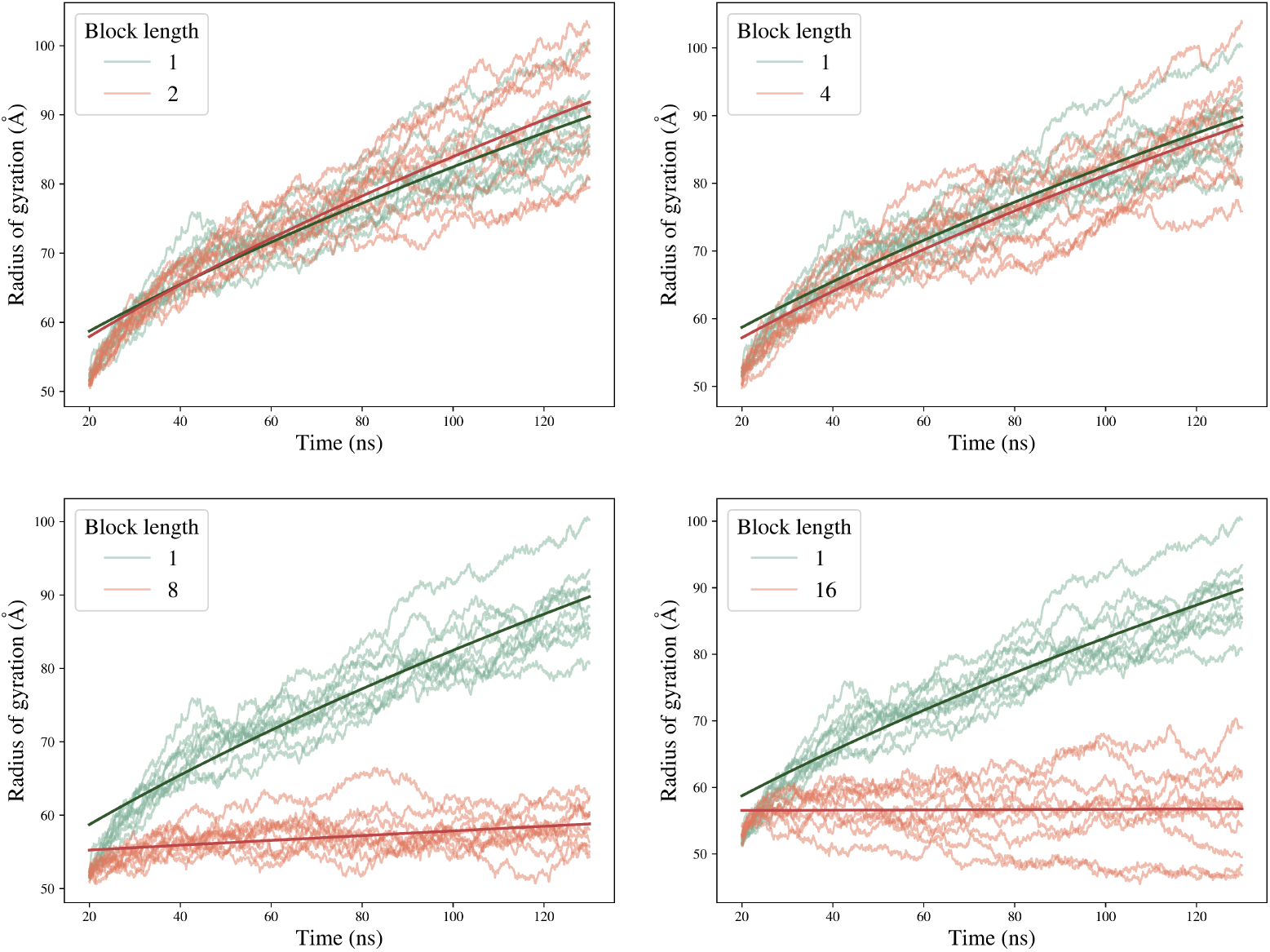
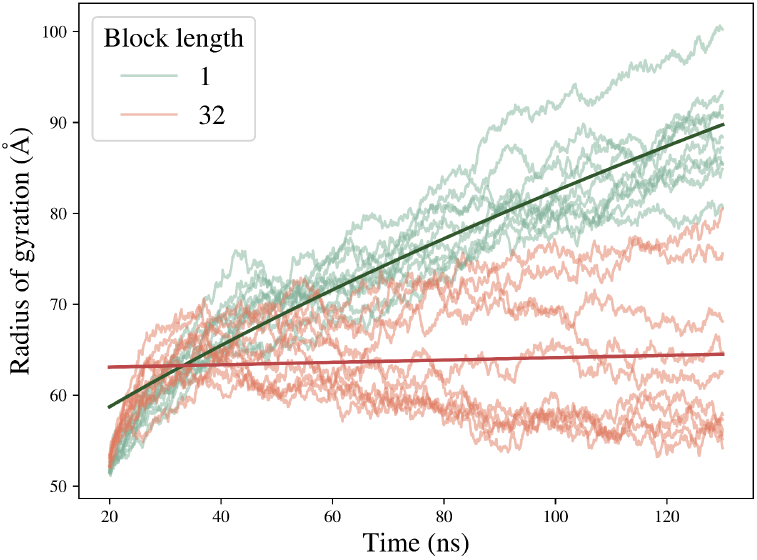
Radii of gyration and their averaged approximations by diffusion equation for varying block lengths compared with such of the homogeneous sequence (block length of one).

### Cluster scaling

Apart from counting contacts, we also used the metric of compactness to characterize grouping of the beads. We found that small side chains are most clustered at a block length of 8. Intermediate blockiness promotes tighter clustering of backbone and spacer beads, whereas higher blockiness excludes spacer residues from the interacting core (Fig. 8). The clustering of stickers becomes most distinguishable for the largest block lengths, indicating that blockiness can be inferred from the clustering of specific bead types (Fig. 9).

**Fig. 8.**
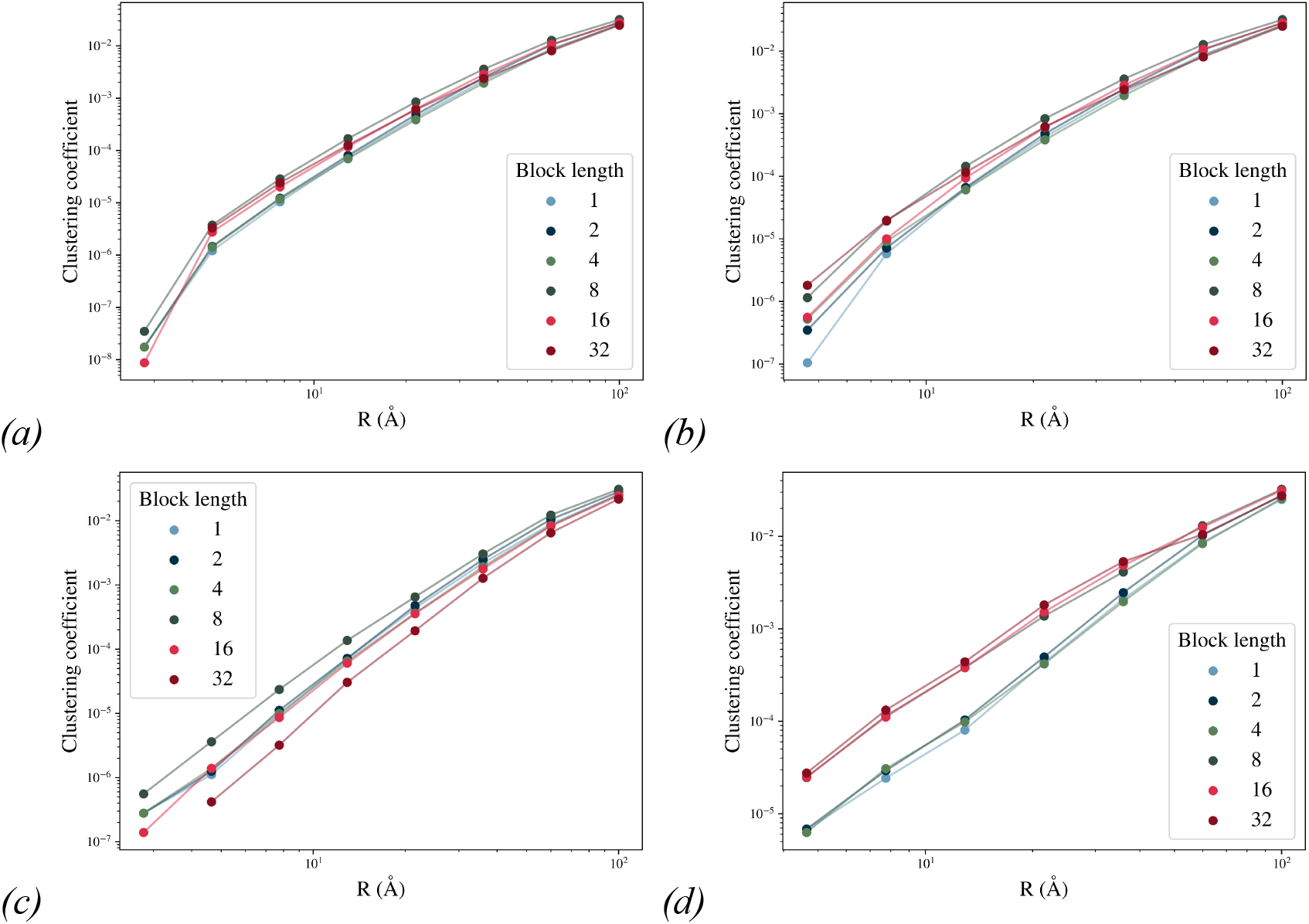
Clustering of MD output structures scales for different blockiness: (a) all beads; (b) backbone; (c) spacer side chain beads; (d) sticker side chain beads.

**Fig. 9.**
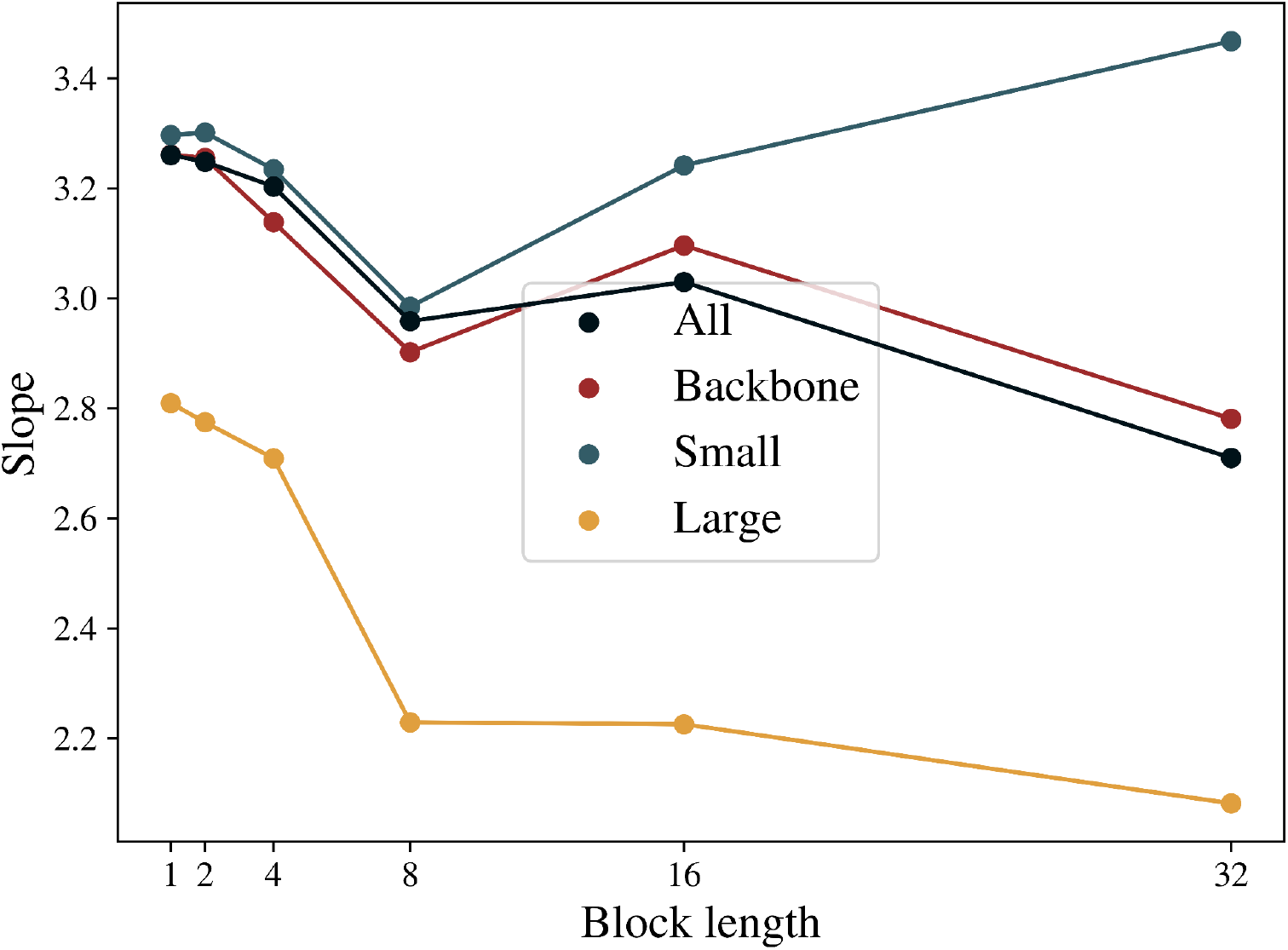
Slopes determined by linear regression of clustering coefficients against distance from Fig. 8; all points except the first were used.

### Sticker-spacer concept and possible architectures of the drivers

Our results demonstrate that the blockwise organization of residues with similar interaction propensities aligns with the stickers-and-spacers approach to LLPS. Driver proteins more frequently exhibit contiguous blocks of bulky residues separated by regions enriched in smaller residues, closely resembling the alternating arrangement of sticker and spacer domains proposed in associative polymer models [Mittag 2022]. This block organisation allows several intermolecular contacts to form simultaneously, thereby a group of residues acts as a binding module. This interpretation is also consistent with previous studies identifying the physicochemical composition of sticker and spacer regions. Sticker domains are typically enriched in bulky aromatic and charged amino acids, such as phenylalanine, tyrosine, tryptophan, arginine, and lysine, whereas spacer regions preferentially contain small, flexible residues including glycine, serine, and other polar amino acid residues [Ji 2026, Bremer 2022]. Our analysis extends these observations by showing that these residues are not only compositionally enriched but are also spatially organized into contiguous blocks. Proteins with variable surface exposure, which facilitates interactions with other proteins and nucleic acids, are more likely to undergo LLPS [Monti 2025]. From a physicochemical perspective, this behaviour can be explained by the combined effects of excluded volume and intermolecular attraction. Residues with larger side chains occupy a greater excluded volume and simultaneously possess stronger van der Waals and hydrophobic interactions, which are commonly represented by the Lennard–Jones potential in coarse-grained molecular models. Multivalent interactions between stickers reduce the entropic penalty associated with network formation and promote the emergence of dense crosslinking regions connected by flexible spacer segments [Ji 2026]. So that blockier sequences form percolated networks at much lower concentrations and formed more stable, less dynamic condensates than sequences in which interacting residues are distributed uniformly. These polymers required substantially higher concentrations, or barely phase-separated at all. Our observations indicate that blockiness determines not only whether aggregation occurs, but also the internal architecture of the condensate. Overall, we show that blockiness of sticker distribution is a fairly required parameter for peptide-like polymer aggregation. Based on our results, we propose several guesses for an effective driver architecture. As shown in Fig. 10, the formation of a stable percolated network with multiple nucleation cores requires at least two sticker domains separated by a sufficiently long spacer in each polypeptide. If the spacer is too short, the polymer preferentially collapses into a single compact aggregate and tends to form rather hexagonal phase than an elastic network.

**Fig. 10.**
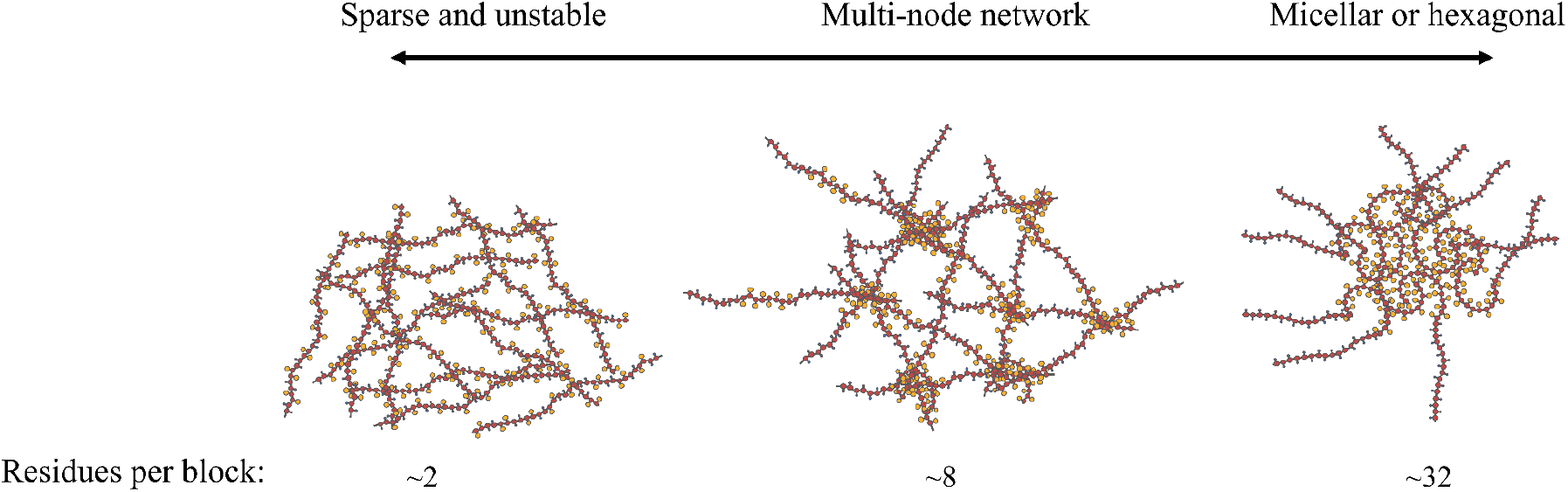
Schematic phase diagram showing the dependence of polymer network topology on block length.

## Conclusions

Taken together, these analyses allow us to assess whether the blockwise organization of stickers contributes to the distinct phase-separation behaviour of driver and client proteins and to evaluate its role in promoting protein aggregation. Residues of similar aggregation propensity tend to be distributed blockwise in the driver proteins and uniformly across the sequences of client proteins. We confirmed this finding by coarse-grained simulations of polymer condensates using polymers with the same composition and varying degrees of residue blockiness. We have shown that more blockwise polymers form a stable aggregate with aggregation cores or hexagonal phase, and polymers with uniformly distributed residues form a flexible percolated network, and there exists blockiness threshold stabilizing polymer networks. Overall, these results identify blockiness of SASA and other properties as a previously unrecognized sequence feature associated with LLPS drivers and further support the importance of residue patterning in the stickers-spacers framework and align with recent developments in molecular grammars.

## Data availability

All scripts for data processing, MD preparation and execution, and output analysis, together with the corresponding outputs, sequences, their asymmetry scores for different window lengths, and obtained statistics, and processed structures ready for visualization, are openly available in the GitHub repository at *https://github.com/ivankovlab/LazyPhase/tree/main/examples/stickers_spacers*.

## Supplementary Data

Representative sequences of LLPS-forming polypeptides and the computed IDR scores for their residues are provided in the ZIP archive accompanying this article.

**Fig. S1.**
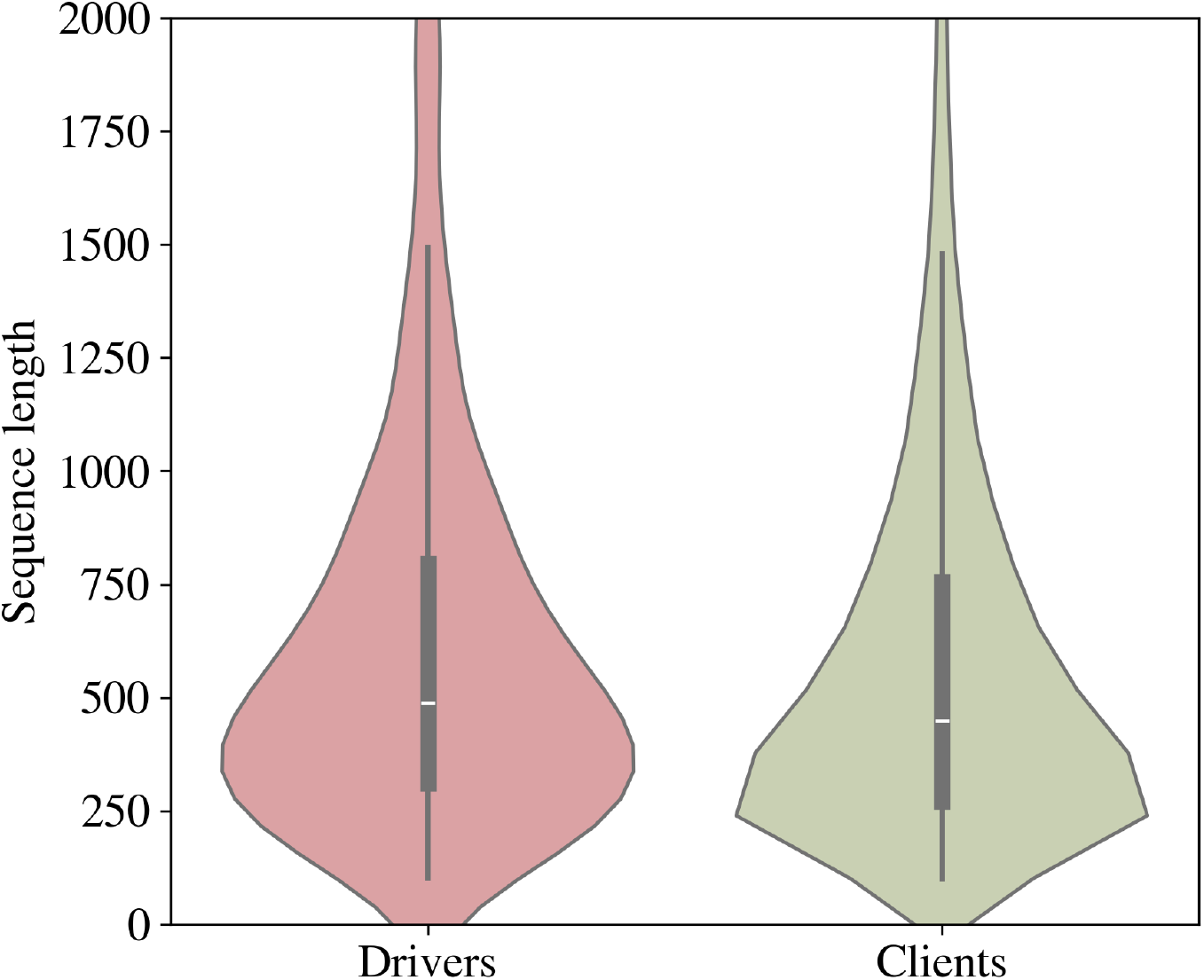
Distributions of driver and client sequence lengths. The mean sequence lengths are 629 and 605 residues, respectively.

